# Theta oscillations show impaired interference detection in the elderly during selective memory retrieval

**DOI:** 10.1101/388595

**Authors:** Catarina S. Ferreira, Maria Jesús Maraver, Simon Hanslmayr, Bajo Teresa

## Abstract

Seemingly effortless tasks, such as recognizing faces and retrieving names, become significantly harder as people get older. These age-related difficulties may be partially due to the concurrent activation of related competitors. However, it remains unclear whether older adults struggle with detecting an early interference signal or with suppressing irrelevant competitors once competition is detected. To investigate this question, we used the retrieval practice paradigm, shown to elicit interference, while recording electrophysiological activity in young and older adults. In two experiments, young participants showed the typical Retrieval Induced Forgetting (RIF) effect whereas the elderly did not. Neurally, young adults were more capable to detect interference than the older, as evidenced by an increase in mid-frontal theta power (~4-8Hz). This efficient interference detection allowed young adults to recruit inhibitory mechanisms that overcome competition, as traced by a theta power reduction across retrieval cycles. No such reduction was found for the elderly, indicating that the lack of an early interference detection signal renders older adults unable to recruit memory selection mechanisms, eliminating RIF.

**AUTHORS NOTE:** This research was supported by the doctoral research grants AP2009-2215 to C.S.F and BES-2013-066842 to M.J.M.; by grants PSI2012-33625; PSI2015-65502-C2-1-P from the Spanish Ministry of Economy and Competiveness, and by the Economic Council of the Andalusian Government P08-HUM-03600-Feder and P12-CTS-2369-Feder to T.B.

**Declaration of interest:** The authors have no conflicts of interest to disclose

## 1. INTRODUCTION

Aging entails a general decline in cognitive functioning, with memory being one of the most affected functions (Park et al., 2002). As people get older, they become more vulnerable to everyday forgetfulness (Cutler and Grams, 1988; Montejo et al., 2012; Ryan, 1992) and perform worse in free recall and recognition tests (Craik, F.; Jennings, 1992; Light, 1991). Furthermore, elders’ memory for names and faces is poorer than that of younger adults (Bahrick, 1984; Cohen & Faulkner, 1986; Maylor, 1990), especially when retrieving face-name associations (Naveh-Benjamin et al., 2004). In fact, recognizing people’s faces and retrieving information about them seems to become particularly demanding with aging, being one of the most commonly reported complaints made by the elderly (Lovelace and Twohig, 1990; Maylor, 1990). Critically, Pike et al. (2012) suggest that deficits in retrieving face-name pairs may help distinguishing between mild cognitive impairment (MCI) and healthy aging, which is especially relevant if we take into account that people with MCI have an increased risk of developing Alzheimer’s disease (Gauthier et al., 2006).

One possible explanation for age related changes was proposed by Hasher and Zacks (1988) in their Inhibitory Deficit Theory (IDT), which posits that cognitive failures related to normal aging are due to a deficit in inhibitory mechanisms. These authors argue that age-related deficits in attention, language or memory, could be due to a common underlying mechanism: a decline in inhibitory function. According to the IDT, older adults do not have the ability to suppress or inhibit unwanted information from entering working memory. Therefore, older people’s naming difficulties could be due to an inability to suppress irrelevant-competing representations making it harder to access and choose the desired information (Lustig, Hasher, & Zacks, 2007). Corroborating this idea, several studies have found age-related impairments when testing participants in inhibitory paradigms (Stop Signal and Go/No go: Bedard et al., 2002; Think/No Think: Anderson, Reinholz, Kuhl, & Mayr, 2011, but see Murray, Anderson, & Kensinger, 2015).

One paradigm commonly used to investigate inhibitory function in selective memory retrieval is the retrieval practice paradigm (Anderson, Bjork, & Bjork, 1994). In this paradigm, participants first study pairs of words belonging to a given category (e.g. FRUIT-Apple; FRUIT-Orange; ANIMAL-Elephant) and are then asked to retrieve half of the words from half of the categories, upon presentation of a cue (e.g. FRUIT–Ap__). The presentation of the category cue (FRUIT) leads to the activation in memory of all previously studied related items, eliciting interference between related competitors (Apple, Orange, Banana…). According to the inhibitory account (Anderson et al., 1994), when interference is detected, a control mechanism is triggered to suppress competing memory representations (Orange) and promote the retrieval of the target memory (Apple). Thus inhibition is required to suppress strong competing responses in order to allow the expression of weaker but more appropriate ones (Levy and Anderson, 2002). Two different memory effects can be observed in a final test. First, a facilitation effect where practiced items (Apple) are recalled significantly better than control items (items that were neither studied nor belong to studied categories, as Elephant). Second, the recall of unpracticed items from practised categories (Orange) is significantly impaired in comparison to control items, an effect named Retrieval Induced Forgetting (RIF, Anderson et al., 1994).

Although alternative explanations have been proposed (Jonker et al., 2013; Mensink and Raaijmakers, 1988) there is overwhelming support for RIF’s inhibitory nature, showing that it occurs due to the inhibition of neural assemblies representing the competitor item (Murayama et al., 2014; Waldhauser, Johansson, & Hanslmayr, 2012; Wimber, Alink, Charest, Kriegeskorte, & Anderson, 2015).

It is important to notice that the inhibitory account implicates at least two mechanisms: i) a mechanism that detects interference, and ii) a mechanism that reduces interference by inhibiting competing memories. Although studies have shown that the behavioural RIF effect is gradually impaired in older adults, and modulated by factors such as age itself (Aslan and Bäuml, 2012; Marful et al., 2015a) or the availability of cognitive resources (Ortega et al., 2012), there are not, to our knowledge, any EEG studies specifically investigating the brain mechanisms underlying age-related changes in RIF. Hence, it remains unclear whether older adults struggle with the detection of an early interference signal or with the suppression of irrelevant competitors. Due to its superb temporal resolution, EEG should allow the dissociation between the two neural signatures mediating RIF (interference detection and inhibition), a difficult goal to achieve when relying purely on behavioral methods, and enable us to identify the source of age-related changes in RIF.

Electrophysiological studies evidence that interference in the retrieval practice paradigm can be traced by mid-frontal theta (~4-8 Hz), with increments in mid-frontal theta power shown when comparing a competitive to a non-competitive condition (Hanslmayr et al., 2010; Staudigl et al., 2010), localized to medial prefrontal brain regions (such as the ACC), and predicting later forgetting (Staudigl et al., 2010). In order to specifically untangle interference and inhibition signals in this paradigm, Ferreira, Marful, Staudigl, Bajo and Hanslmayr (2014) presented a category cue (e.g. Actor) and a retrieval specific cue (the face of a specific actor) separated in time. Mid-frontal theta oscillations were attributed specifically to interference detection, by showing an increase in theta power in a competitive (vs. a non-competitive) condition, upon presentation of the category cue, when competing items become active in memory. Theta power decreased in the competitive condition from the presentation of the category cue to the presentation of the retrieval cue, reflecting a reduction in interference or its resolution, which correlated with later forgetting.

This paradigm, in combination with electrophysiology, is therefore ideally suited to reveal the mechanism potentially impaired in older adults: interference detection or its resolution (i.e. inhibition). In two consecutive experiments (using faces and semantic material) we employ a procedure similar to that of Ferreira et al. (2014b). We compare the neural correlates of RIF throughout subsequent cycles of retrieval practice between age groups. We assess theta oscillations upon presentation of the category cue, to disentangle interference detection (first presentation of the cue, when interference should be at its highest level) and inhibition or interference resolution (difference between first and last presentation of the cue).

If the elders’ difficulties in naming faces are due to poor interference detection, we would expect theta power upon presentation of the first cue to be lower for older compared to younger participants (Figure 1A), as supported by studies showing that low forgetters exhibit lower levels of theta than high forgetters (Staudigl et al., 2010), as well as lower ACC activity (Kuhl et al., 2007) on the first retrieval practice cycle. In this case, theta power should remain constant across retrieval cycles, since inhibitory mechanisms should not be called into play unless interference is detected (Anderson, Bjork, & Bjork, 2000). If, on the other hand, the problem lies on interference resolution, theta power upon presentation of the first retrieval cue should be equivalent between young and older participants, and again, should remain constant (at high levels, in this case) across cycles for the elderly, as they would not be able to engage the necessary mechanisms to solve interference amongst stimuli.

**Figure 1:**
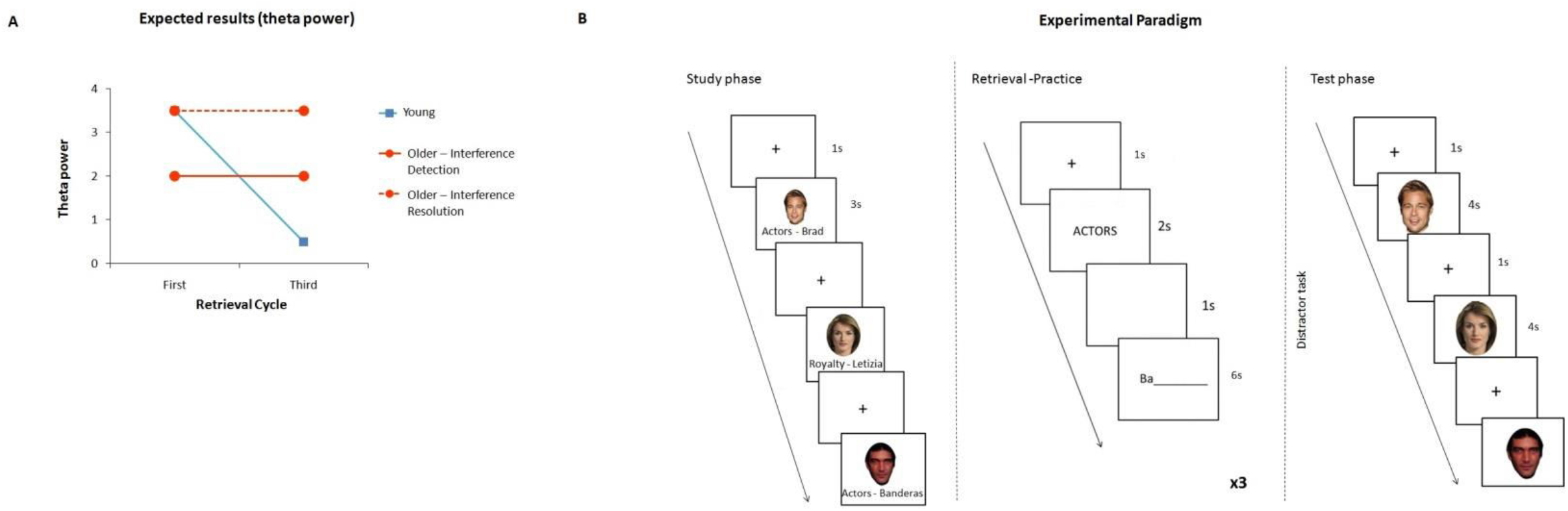
**A)** Expected neural results. The blue line represents the expected results in theta power for the young participants, replicating previous studies. The orange lines represent the expected results for the elderly. The solid line represents the expected results if older participants suffer from a deficit in interference detection, whereas the dashed line depicts what would be expected if the elders’ struggle is in solving interference. **B)** Depiction of the experimental paradigm.

## 2. EXPERIMENT 1

### 2.1 Methods

#### 2.1.1 Participants

24 students from the University of Granada (17 female; *M_age_*=24.70; *SD*=5.56) and 24 older adults (10 female; *M_age_*=68.38; *SD*=5.18; range 60-79) participated in this study^1^. Older participants were recruited from an association for retired people. These participants were highly educated (*M_scholarity(years)_*=13.31; *SD*=3.25) and going through a process of normal healthy aging (Mini Mental State Exam (Lobo, Ezquerra, Gómez Burgada, Sala, & Seva Díaz, 1979) mean score = 28.65/30, *SD*=1.37) There were no significant working memory differences between the two groups, as measured by the digits span test from the Wechsler Adult Intelligence Scale (WAIS III; *M_young_*= 15.13, *SD_young_*=2.82; *M_older_*=13.58, *SD_older_*=2.99; *p*>.05).

All participants were Spanish or had been living in Spain for at least 15 years and all reported normal or corrected-to-normal vision. Participants were given all the information about the study and signed an informed consent prior to its start. All subjects received course-credits or a monetary reward for their participation in the study.

For the EEG analysis, four of the older participants were excluded due to excessive movement during the task leading to poor EEG data quality.

#### 2.1.2 Material

A total of sixty-four pictures were used in this experiment, all of famous people. Faces were divided into eight occupational categories (male: actors, politicians, football players, writers, and TV hosts; female: singers, royalty members, and tabloid stars). These materials had been used in previous experiments (Ferreira et al., 2014a, 2014b) and were originally chosen from a pilot study that served the purpose of evaluating each item’s familiarity. Pictures were selected so that they had the highest familiarity values, provided they did not share the first two letters of the corresponding name. Six additional exemplars were chosen as filler items: 3 radio personalities and 3 bull-fighters. These were used to control for primacy and recency effects and were not taken into account in any of the analyses.

Pictures were presented in colour (5.19cm×6.99 cm) against a white background. In order to standardize them, an oval template was applied around each picture (see Young, Ellis, Flude, McWeeny, & Hay, 1986). All faces displayed a neutral to mildly positive expression. Eight counterbalance versions were created so that all faces were seen in all conditions across participants.

### 2.2 Procedure

The experiment consisted of a version of the retrieval practice paradigm (Figure 1B), comprising a study phase, a retrieval practice phase and a final test.

#### 2.2.1 Study phase

The experiment started with a study phase, where participants were shown the 64 critical faces sequentially. Presentation was randomized except that the first and last three faces were always filler items, to account for primacy and recency effects. After a 1000ms fixation cross, a face appeared on the screen for 4000ms with its respective name and profession written below (e.g. Actors–Banderas). The participants’ task consisted in pressing a number from 1 to 5 on the keyboard to rate how familiar they were with the face presented on the screen (1-not known at all; 5-very well known). This was done not only to control for possible differences in item familiarity between older and younger participants, but also to keep participants engaged and ensure they attended to and processed the stimuli. Subjects were instructed to pay close attention not only to the faces but also to their names and professions since they would be asked about them in the next phase.

#### 2.2.2 Retrieval practice phase

During this phase, which occurred right after study, participants were asked to retrieve half of the exemplars from six of the eight categories. Participants first saw a jittered fixation cross (1000-1500ms) followed by the category cue (e.g. Actors) for 2000ms, a blank screen (500ms) and a specific face (2500ms). Then a red question mark appeared on the screen and participants were instructed to give their response (name the person they had just seen on screen) at that moment. Participants were explicitly asked to refrain from responding until the question mark was presented on screen, to avoid speech artefacts. Faces were presented in a pseudo-random order, so that a whole set would be presented before repeating itself. As in the study phase, the first and last faces were filler items used to control for primacy and recency effects.

Crucially, there were three cycles of retrieval practice, that is, each of the 24 critical faces used during this part of the experiment was repeated three times, in order to allow comparisons between first and third cycles, similarly to what has been done in previous studies (Kuhl et al., 2007; Staudigl et al., 2010;Wimber, Rutschmann, Greenlee, & Bäuml, 2009).

#### 2.2.3 Test phase

A 5 minute distracter task followed retrieval practice (the digits span test from the WAIS III). Thereafter, a final memory test took place, where each studied face was presented again for naming. After a fixation cross (1000ms), a face appeared on the screen for 3000ms and participants were asked to retrieve the corresponding name as soon as possible. The order of presentation was randomized, such that all unpractised items and half of the control items were presented first, followed by practised items and the other half of the baseline-items. This was done to prevent possible confounds regarding the forgetting effect, as retrieval of practised items first could block access to the unpractised ones (blocking effect: Kohler & McGeoch, 1943; Mensink & Raaijmakers, 1988).

### 2.3 EEG Recording

The EEG was recorded from 64 scalp electrodes, mounted on an elastic cap, on a standard 10-20 system. Four additional electrodes were used to control for eye movements: two set above and below the left eye (controlling for vertical movement) and another two set at the outer side of each eye, to control for horizontal movement.

Continuous activity was recorded using Neuroscan Synamps2 amplifiers (El Paso, TX) and was first recorded using a midline electrode (half-way between Cz and CPz) as reference. The data was then re-referenced offline against a common average reference. Each channel was amplified with a band pass of 0.01-100Hz and digitized at a 500Hz sampling rate. Impedances were kept below 5kΩ.

Prior to analysing the data, a high-pass filter (at 1Hz) was applied and artefacts (such as eye movements and EKG) were removed using independent component analysis (ICA). Remaining artefacts after ICA were manually removed by carefully inspecting the data.

### 2.4 EEG pre-processing

For EEG analyses we used the Fieldtrip toolbox (Oostenveld et al., 2011) on Matlab (The MathWorks, Munich, Germany). The EEG data were cut into segments ranging from - 2000ms before stimulus presentation to 4000ms after, around both the category cue and the retrieval specific cue (i.e. the face; first, second, and third cycle in both cases). These large segments were chosen to avoid filter artefacts after wavelet transformation at the beginning and end of each period. Data analysis was restricted to a smaller time window from -500ms to 2000ms.

### 2.5 Analyses of Oscillatory Power

For time-frequency analyses, a Morlet wavelet transformation (7 cycles) was applied to the data. Data were filtered in a frequency range from 1-30 Hz and exported in bins of 50ms and 1Hz. As in previous experiments (Ferreira et al., 2014b), power changes were calculated in relation to a prestimulus baseline (from -500 to 0ms before category cue onset).

Given our a-priori hypotheses, analyses were restricted to the theta frequency range (4-8Hz) and to the time window around cue presentation. A region of interest analysis was applied on a set of 9 fronto-central electrodes (Fcz, F1, Fz, F2, Fc1, Fc2, C1, Cz, C2) based on our previous study (Ferreira et al., 2014b) and on a plethora of other studies showing that mid-frontal theta oscillations are typically recorded at these locations (Cohen, 2014). We believe that restricting our analyses to this ROI, allows for more specific interpretations of the results, since we have very clear a-priori hypotheses about what mid-frontal theta should be reflecting (for unrestricted analyses, however, see Supplementary Figure 1). Power differences over this ROI were used to define the exact time-frequency windows for subsequent analyses.

Since the aim of this study was to assess differences between young and older adults, our first step was to compute group differences. We first looked at differences in theta power upon presentation of the first category cue, as an index of initial levels of interference detection, and then performed an interaction analysis (cue cycle 1 minus cue cycle 3 × age group). Differences in theta power upon presentation of the cue on the first cycle minus presentation of the cue on the third one were calculated for each participant, at the aforementioned mid-frontal ROI. These differences were then subjected to an independent samples *t*-test, comparing the two age groups. Note that although age-related anatomical differences, such as skull thickness, could have an effect on absolute power, they should not affect these relative measures.

Analyses of oscillatory power upon face presentation were performed in a similar fashion to the category cue analyses, although differences were computed for all electrodes, rather than for a particular ROI. These are not reported since they yielded no significant results.

In order to control for multiple comparisons, Monte Carlo randomization was used (see details on this method in Maris & Oostenveld, 2007). From this procedure, clusters of electrodes that significantly differed from one cycle to the other were obtained (*p_corr_*<.05).

Planned comparisons were then made for each group (young and older) separately, comparing first cue and face presentations minus third, over the time and frequency windows significant in the interaction analysis.

### 2.6 Results

#### 2.6.1 Behavioural Results

Familiarity ratings at study did not differ overall between groups (*M_young_*=4.00; *SD_young_*=.78; *M_older_*=4.10; *SD_older_*=.53; *p*>.05). Dividing the items according to occupational category revealed that older adults were more familiar with writers than the younger adults (*M_young_*=3.00; *SD_young_*=.82; *M_older_*=4.30; *SD_older_*=.39; *t*(14)=-3.84, *p*>.01), whereas the opposite was true for football players (*M_young_*=4.70; *SD_young_*=.15; *M_older_*=3.90; *SD_older_*=.26; *t*(14)=7.74, *p*>.001). No other categories differed between groups.

Mean recall during the retrieval practice phase did not differ either between young and older adults (*M_young_*=.76; *SD_young_*=.16; *M_older_*=.67; *SD_older_*=.18; *p*>.05).

Two 2×2 repeated measures ANOVA were conducted to assess forgetting and facilitation effects separately on the final memory test. Post-hoc analyses were conducted for each group, using 1-tailed paired-sample *t*-test. A summary of the descriptive statistics is detailed in Table 1.

**Table 1:** Summary of the behavioural descriptive statistics from Experiment 1 (Face material).

|  | Young |  | Old |  | Group effect |  |
| --- | --- | --- | --- | --- | --- | --- |
|  | *M* | *SD* | *M* | *SD* | *p* | *Cohen's d* |
| Age | 24.70 | 5.56 | 68.38 | 5.18 | 0.00 | -8.13 |
| Working Memory<br>(Digits WAIS-III) | 15.13 | 2.82 | 13.58 | 2.99 | 0.07 | 0.53 |
| Retrieval Practice<br>(overall recall) | 0.76 | 0.16 | 0.67 | 0.18 | 0.10 | 0.53 |
| Final memory test<br>(overall recall) | 0.66 | 0.20 | 0.51 | 0.15 | 0.00 | 0.85 |
| Control (Nr<sub>p</sub>) | 0.65 | 0.25 | 0.46 | 0.22 | 0.01 | 0.81 |
| Unpractised (Rp<sub>-</sub>) | 0.58 | 0.22 | 0.43 | 0.15 | 0.00 | 0.80 |
| *Forgetting* | 0.07 | 0.17 | 0.03 | 0.20 | 0.51 | 0.21 |
| Practised (Rp<sub>+</sub>) | 0.75 | 0.17 | 0.64 | 0.17 | 0.03 | 0.65 |
| *Facilitation* | 0.10 | 0.16 | 0.18 | 0.20 | 0.11 | -0.44 |

##### 2.6.1.1 Forgetting

The results of the ANOVA type of item (unpractised vs. control) × group (younger vs. older) showed a significant effect of group [*F*(1,46)=10.03,*p*<.01, *η*_*p*_^2^=.18], where younger participants’ overall proportion of recall was higher than older participants’ (see Table 1). A marginally significant effect of item type [*F*(1,46)=3.64, *p*=.06, *η*_*p*_^2^=.07] was also found, with lower mean recall of unpractised items than of controls. Although the interaction between age group and item type did not reach significance [*F*(1,46)<1), n.s.], planned contrasts were conducted nonetheless, given that we had very specific hypotheses of what to expect. These revealed that whereas the difference between unpractised and control items was significant for the younger adults [*t*(23)=-1.97,*p*<.05], such difference was not significant for older participants.

##### 2.6.1.2 Facilitation

Regarding the facilitation effect, no significant item type (practised vs. control) × age group interaction was found [*F*(1,46)=2.66, *n.s*.]. However, there was a significant main effect of age group [*F*(1,46)=7.92, *p*<.01, *η_p_*^2^=.78], according to which younger adults recalled more items overall than older adults did. Moreover, item type also reached statistical significance [*F*(1,46)=30.26,*p*<.001, *η_p_*^2^=.39]. Recall of practised items was significantly better than recall of control items. Both young [*t*(23)=3.1,*p*<.01] and older [*t*(23)=4.6, *p*<.001] participants recalled practised items significantly better than baseline (see Table 1).

#### 2.6.2 Theta Power Results

##### 2.6.2.1 Young vs. Old: Cue 1 and Cue 1 vs. Cue 3

Differences in theta power upon presentation of the cue on the first and third cycles were computed for each participant in the young and older group. We first report the analysis for the first cycle (Cue 1; interference index) and then the difference between the first and third cycles (Cue 1 vs. 3; interference resolution).

For the first cue presentation a significant difference in theta power was found between younger and older adults (*p*_corr_<.01), such that younger adults showed greater theta power (7-8 Hz) over the mid-frontal ROI, in a time window ranging from 0 to 500 ms (Figure 2A).

**Figure 2:**
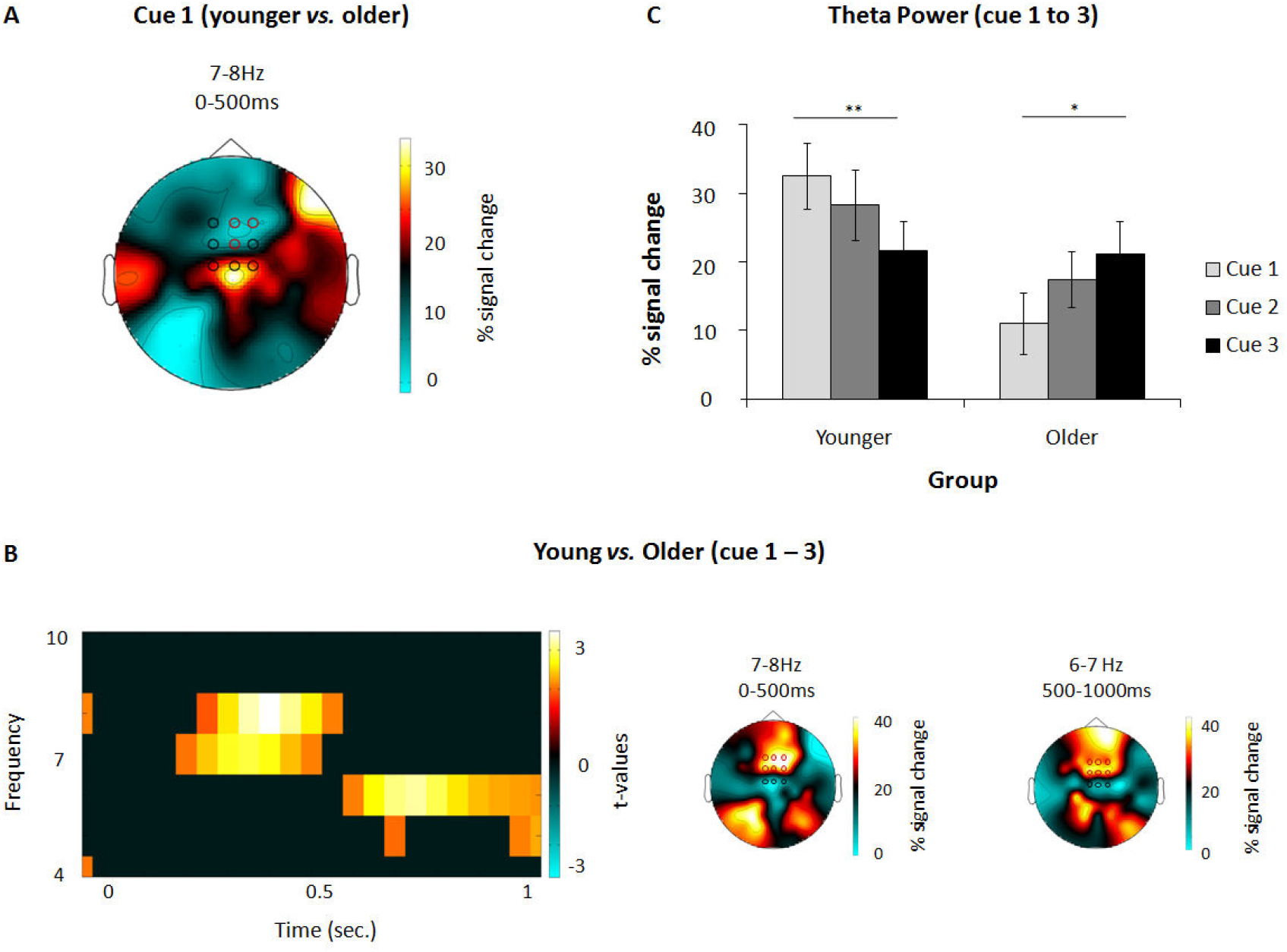
**A)** Topography depicting differences in activity between younger and older participants, upon presentation of the category cue. The ROI electrodes are depicted in a black circle. Significant ones are highlighted in red. **B)** Interaction analysis: differences between younger (cue1 – cue3) and older adults (cue1 – cue3). On the left, the time-frequency plot shows the significant time-frequency windows used for subsequent analyses and the topographies on the right show the distribution of these effects. All the analyses leading to plots A) and B) where conducted over a central ROI comprising 9 mid-frontal electrodes (depicted in black circles), and electrodes that showed significant differences can be seen in red circles. **C)** The graph bars show the percentage signal change in theta power (7-8Hz), from 0 to 500ms upon presentation of the category cue in each cycle for young (left) and older (right) participants. Note how theta power decreases across retrieval cycles for the younger participants but shows an opposite patter for the elderly (\**p*<.05; \*\**p*<.01).

The interaction analysis (first cue minus third cue × age group) yielded a significant effect in two different time-frequency windows, over the mid-frontal ROI. In both windows theta power was higher for younger compared to older adults (see Figure 2B). The first time-frequency window ranged from 7-8 Hz in the first 500ms upon stimulus onset (*p*_corr_<.001); the second significant window ranged from 6-7 Hz at 500 to 1000ms (*p*_corr_<.01). Follow-up comparisons on these two effects are described next.

##### 2.6.2.2 Young adults: Cue 1 vs. Cue 3

As expected, young adults showed a significant theta power decrease upon cue presentation from the first to the third retrieval practice at the sensor level, both from 7-8Hz during the first 500ms (*p*_corr_<.01) and from 500 to 1000ms, at a frequency range from 6-7 Hz (*p*_corr_<.01). In order to get a clearer picture of how theta power progresses from one cycle to another, we extracted theta power values upon presentation of the category cue, in the two significant time windows and over the 9 electrode ROI, for the three cycles. The results of this analysis indicated that in both time-frequency windows, theta power gradually decreased across cycles. For both time-windows, the difference between first and third cue reached statistical significance (first time-window, depicted in Figure 2C: *t*(23)= 2.81, *p*<.01; second time-window: *t*(23)= 2.47, *p*<. 05].

##### 2.6.2.3 Older adults: Cue 1 vs. Cue 3

Older adults also showed a significant modulation of mid-frontal theta power across retrieval cycles in the first time window, from 0 to 500ms and from 7-8Hz (*p*_corr_<.05).However, in stark contrast to the younger adults, theta power increased from the first to the third category cue. No significant difference emerged in the second time window (500-1000ms; 6-7Hz; all *p*_corr_>.05). We again extracted theta power values upon presentation of the cue for each retrieval practice cycle. As expected from the results found in the interaction analysis, older people seem to have, overall, lower levels of theta power compared to the younger. Theta power increased numerically from the first to the second and third retrieval cycles, with differences between first and third cue reaching statistical significance on the first time-frequency window [*t*(19)=-1.89, *p*<.05, Figure 2C].

## 3. EXPERIMENT 2

The results from Experiment 1 suggest that whereas in younger adults interference detection triggers the memory selection mechanisms necessary to suppress competing representations, these mechanisms might be impaired in the elderly. This suggests older people might perform worse in inhibitory tasks due to the lack of an early interference signal that would be responsible for triggering inhibitory mechanisms that solve competition.

An alternative explanation is that older adults are simply not processing the category cue, which is supposed to elicit or boost interference, to the same extent that young adults do. This idea is supported by studies showing impaired context processing in older adults (Braver et al., 2001; Rajah et al., 2010; Rush et al., 2006). Older participants might simply not be processing the context (the category cue), due to this impairment, combined with the fact that the cue is not essential to perform the task correctly (participants can still recall the name “Banderas” upon face presentation, regardless of whether or not they read the cue “Actors” beforehand).

To rule out these alternative accounts, we ran a second experiment, using pairs of words instead of faces. We presented a category cue (FRUIT) followed by the word stem of an exemplar of that category (Ap___ to retrieve Apple). Using semantic material forces participants to focus on the category cue, since in order to successfully retrieve the target “Apple” one needs more information than the word’s stem. This necessary information is given by the category cue (FRUIT). Therefore participants cannot afford to ignore the cue in order to retrieve the correct target.

If the results from Experiment 1 were due to impairment in context processing or to the use of different strategies by the elderly, by forcing them to focus on the category cue, we should find results (both at the behavioural and neural level) more in line with those found in the younger adults. If, however, older adults are indeed less capable of detecting competition, then we should be able to replicate the results from the first Experiment.

### 3.1 Methods

#### 3.1.1 Participants

24 students from the University of Granada (17 female; *M_age_*=21.13; *SD*=3.45) and 24 older adults (8 female; *M_age_*=64.74; *SD*=3.45; range 60-75) participated in this study. Older participants were recruited from an advert published in a local newspaper and on the University of Granada webpage. Inclusion criteria specifically stated that older adults should have a minimum of 12 years of education. Mean years of education for this sample was of 15.46 (*SD*=2.25). As in Experiment 1, participants completed the Mini Mental State Exam (MMSE; Lobo et al., 1979), scoring 28.1/30 (*SD*=0.98). No differences were found between groups as to working memory capacity, measured by the digits span test from the WAIS III (*M_young_*=15.88; *SD_young_*=2.58; *M_older_*=14.53; *SD_older_*=2.98; *p*>.05).

All participants were Spanish or had been living in Spain for at least 15 years and were thus native or very fluent speakers. All reported normal or corrected-to-normal vision. Participants were given all the information about the study and signed an informed consent prior to its beginning. Young participants received course-credits and older adults were monetarily rewarded for their participation in the study.

#### 3.1.2 Material

A total of 64 target words plus six fillers were used. The words belonged to eight different categories (animals, fruits, tools, vehicles, insects, trees, clothes and furniture) with eight exemplars each. Filler items belonged to two extra categories (beverages and toys, with three exemplars each).

Within the same category, no items shared the first two letters. Moreover, in order to maximize competition between items, within each category four items were highly representative of their categories, while other four were poor representatives. The poor representatives were used as practised items and their baseline, whereas highly representative words were used as unpractised ones and their respective baseline. This manipulation is thought to boost interference, since the more representative of its category an item is, the more it will compete with the to-be retrieved ones (Anderson et al., 1994; Anderson, 2003).

Indices of frequency and rank for each item respective to its category were taken from Marful, Díez, & Fernandez (2015), using the NIPE database (Norms and Indices for Experimental Psychology; Díez, Fernandez, & Alonso, 2014). Mean frequency was of 4.70 (*SD*=6.20) for practice items and 208.3 (*SD*=52.80) for competitors. Rank scores were on average 8.5 (*SD*=2.20) and 4.4 (*SD*=1.50) for practice and competitors respectively.

The words were presented in the centre of the screen in a black font (Courier New, 18 pts) on a white background. Category cues were always presented in uppercase letters, whereas the specific items and their stems were presented in a capitalized fashion.

### 3.2 Procedure

Experiment 2 followed the procedure of Experiment 1 as closely as possible. The three phases of the retrieval practice paradigm were maintained, the only difference being that instead of seeing faces together with their respective name and occupation, participants saw a category cue (e.g. FRUIT) together with an exemplar of that same category (e.g. Apple).

### 3.3 EEG Recording, pre-processing and Analyses of Oscillatory Power

The EEG recording and pre-processing were done using the same parameters and equipment as in Experiment 1. Analyses of oscillatory power were conducted in the same fashion as Experiment 1.

### 3.4 Results

#### 3.4.1 Behavioural Results

Descriptive behavioural statistics for Experiment 2 are summarized in Table 2. For the intermediate retrieval practice phase, no differences in mean recall were found for the two age groups (*p*>.05).

**Table 2:** Summary of the behavioural descriptive statistics from Experiment 2 (Semantic material).

|  | Young |  | Old |  | Group effect |  |
| --- | --- | --- | --- | --- | --- | --- |
|  | *M* | *SD* | *M* | *SD* | *p* | *Cohen's d* |
| Age | 21.13 | 3.52 | 64.74 | 3.51 | 0.00 | -12.42 |
| Working Memory<br>(Digits WAIS-III) | 15.88 | 2.58 | 14.83 | 2.98 | 0.20 | 0.38 |
| Retrieval Practice<br>(overall recall) | 0.57 | 0.13 | 0.58 | 0.11 | 0.72 | -0.11 |
| Final memory test<br>(overall recall) | 0.58 | 0.10 | 0.57 | 0.08 | 0.79 | 0.09 |
| Control (Nrp High) | 0.79 | 0.14 | 0.68 | 0.20 | 0.05 | 0.59 |
| Unpractised (Rp-) | 0.67 | 0.15 | 0.70 | 0.10 | 0.40 | -0.25 |
| *Forgetting* | 0.12 | 0.18 | -0.02 | 0.25 | 0.04 | 0.62 |
| Control (Nrp Low) | 0.29 | 0.17 | 0.34 | 0.22 | 0.39 | -0.26 |
| Practised (Rp+) | 0.58 | 0.14 | 0.57 | 0.10 | 0.76 | 0.09 |
| *Facilitation* | 0.29 | 0.20 | 0.23 | 0.25 | 0.35 | 0.28 |

Behavioural analyses were conducted in the same way than Experiment 1, but note that in this experiment practised and unpractised items were contrasted against their own control items, drawn from equally low or high representativeness of their category, respectively.

##### 3.4.1.1 Forgetting

The ANOVA conducted to assess forgetting (type of item × age group) revealed no significant effect of item type [*F*(1,45)=2.51, *p*>.05] or age group [*F*(1,45)=1.69, *p*>.05], but did yield a significant item × group interaction [*F*(1,45)=4.88, *p*<.05, *η_p_*^2^=.10].

Post-hoc analyses showed a significant difference between unpractised items and their respective controls (see Table 2) for the younger adults [*t*(23)=-3.19, *p*<.01]. No such difference was found however in the older group.

##### 3.4.1.2 Facilitation

Concerning the facilitation effect (see Table 2), a significant main effect of item type was obtained [*F*(1,46)=61.72,*p*<.001, *η_p_*^2^=.57], according to which practised items were recalled significantly better than their controls. No significant main effect for group [*F*(1,46) <1, *p*>.05] or item × group interaction [*F*(1,46)=1.20, *p*>.05] were obtained. Post-hoc analyses revealed that facilitation effects were present in both groups [young: *t*(23)=7.23, *p*<.001; older: *t*(23)=4.30, *p*<.001].

#### 3.4.2 Theta Power Results

##### 3.4.2.1 Young vs. Old: Cue 1 and Cue vs. Cue 3

Differences in theta power upon presentation of the cue on the first and third cycles were computed for each participant in the young and older group. As in Experiment 1, we first report the analysis for the first cycle (Cue 1; interference detection index) and then the difference between the first and third cycles (Cue 1 vs. 3; interference resolution).

For the first cue presentation a significant difference in theta power was found between younger and older adults (*p*_corr_<.02), such that younger adults showed greater theta power (7-8 Hz) compared to older adults over frontal and parietal areas, in a time window ranging from 200 to 500ms (Figure 3A).

**Figure 3:**
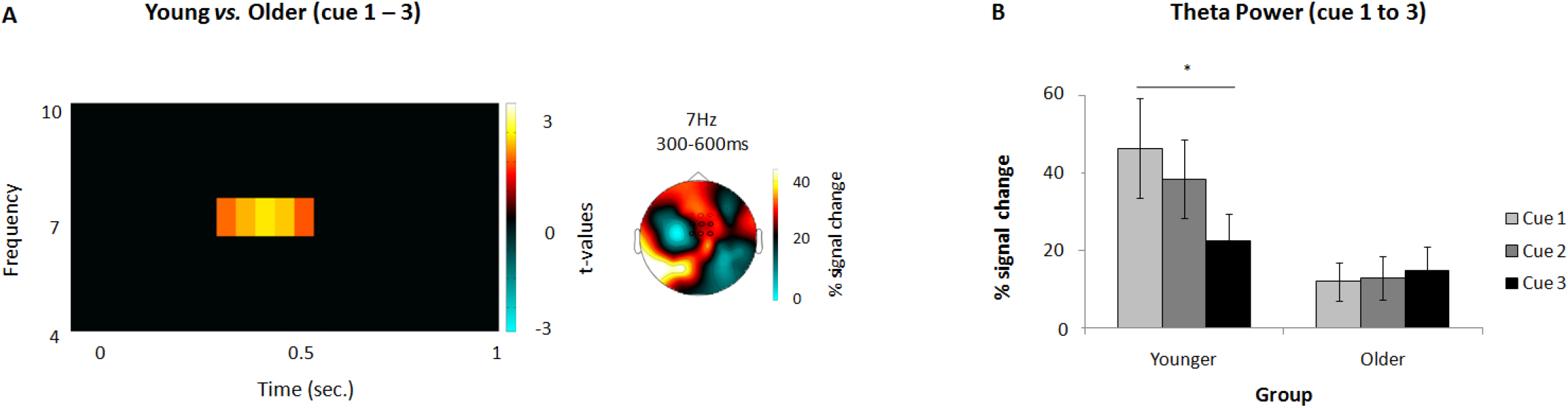
**A)** Interaction analysis: differences between younger (cue1 – cue3) and older adults (cue1 – cue3). The time-frequency plot on the left shows the significant time-frequency window (over a central ROI comprising 9 mid-frontal electrodes, depicted in black circles) used for subsequent analyses and the topography on the right shows the distribution of this effect. Electrodes that showed significant differences can be seen in the red coloured circles. **B)** Percentage signal changes in theta power (7Hz), from 300 to 500ms upon presentation of the category cue in each cycle for young (left) and older (right) participants. Note how theta power decreases across retrieval cycles for the younger participants but not for the elderly (\**p*<.05; \*\**p*<.01).

The interaction analysis (first cue minus third cue × age group) yielded a significant difference over a time window ranging from 200 to 500ms, at 7Hz (*p*_corr_<.02; see Figure 4A). Over the mid-frontal ROI, theta power was higher for younger compared to older adults. Planned comparisons on this effect are described next.

##### 3.4.2.2 Young: Cue 1 vs. Cue 3

For young adults, mid-frontal theta power decreased at 7 Hz upon cue presentation from the first to the third retrieval practice (*p*_corr_<.02)from 200 to 500ms. Replicating the results from the previous experiment, theta power gradually decreased from the first to the third cycle (Figure 3B). The differences in theta power between the first and third cue were statistically significant [*t*(23)= 2.11,*p*<. 05].

##### 3.4.2.3 Old: Cue vs. Cue 3

For older participants, no significant differences were found between the first and the third cue.

## 4. GENERAL DISCUSSION

Across two experiments using different retrieval practice paradigm (Anderson, Bjork & Bjork, 2004) with different materials, younger adults exhibited the typical RIF effect (unpractised items recalled below baseline), while this effect was absent in the elder group.

We argue that whereas younger adults inhibited competing items to promote the correct recall of targets, older adults were not capable of suppressing these irrelevant memories. Importantly, older adults did benefit equally from repeated retrieval, given that in both experiments there were no differences in the average recall during the retrieval practice and that the facilitation effects were similar in younger and older adults.

To rule alternative explanations, Experiment 2 used semantic material, forcing participants to focus on and process the category cue. Crucially, behavioural RIF was still absent in the older group. Note that alternative accounts of RIF, such as blocking (Mensink and Raaijmakers, 1988) or contextual theories (Jonker et al., 2013) cannot fully explain our results either.

This leaves open three possibilities: i) the absence of RIF is due to a poor detection of interference (that is, participants do not detect interference and consequently do not trigger the necessary suppression mechanisms); ii) the lack of RIF occurs due to an inhibitory deficit (participants do detect interference but suffer from an inhibitory deficit that does not allow them to overcome this interference) or iii) both of these processes underlie impaired RIF in the elderly.

Since this is a difficult question to address on a purely behavioural level, we focused on mid-frontal theta power as a proxy for interference detection (Cohen, 2014). Interference should be highest during the first cycle (Kuhl et al., 2007; Staudigl et al., 2010). Indeed, our results show that mid-frontal theta power was higher for younger than older participants, in a time-frequency window ranging from 7-8Hz and 0 to 500ms, which is in good agreement with previous results (Ferreira et al., 2014b; Hanslmayr et al., 2010; Staudigl et al., 2010).

The fact that older adults showed lower levels of theta power than younger participants parallels results showing that lower forgetters exhibit less theta power during the first retrieval cycle (Staudigl et al., 2010) and less ACC activity (Kuhl et al., 2007) than high forgetters. Moreover, it is well known that prefrontal structures suffer from aging to a great extent, and age-related atrophy in frontal lobes has been closely linked to a decrement in cognitive functioning (Nielson et al., 2002; Raz, 2000). Cummins and Finnigan (2007) found altered frontal/ACC theta power in older adults and Pardo et al. (2007) showed a decrease of glucose uptake with aging in the ACC, which correlated with a decline in cognitive performance. Thus, the fact that older participants show less theta power upon cue presentation indicates they do not efficiently engage the brain mechanism in charge of detecting and reacting to interference, which is in line with studies showing this population is more susceptible to interference (Friedman and Castel, 2013; Solesio-Jofre et al., 2012; Tays et al., 2008). Altered ACC function, along with inefficient connection between fronto-parietal regions, within a neural network relevant to perform a memory task (as shown in Pinal, Zurrón, Díaz and Sauseng (2015)), could potentially underlie the reduced mid frontal theta power that older participants show across the two studies presented here.

A significant reduction in theta power from the first to the third category cue was found in the younger adults group, in both experiments. Theta power decreased gradually from one cycle to the next arguably reflecting the successful down-regulation of interference, a marker of how successful inhibition was (Kuhl et al., 2007; Staudigl et al., 2010; Wimber et al., 2011). The more effective detection of interference by younger adults allowed them to trigger the necessary inhibitory mechanisms, which suppress competing items and to therefore resolve interference. This successful suppression of competing items (Waldhauser et al., 2012;Wimber et al., 2015), promotes the correct recall of the sought after target memories in younger adults.

Remarkably, no such theta power reduction was found for the older participants’ group. Our results show that theta power was either constant across retrieval cycles (Experiment 2), or even went in the opposite direction, with theta power increasing from the first to third retrieval cycle (Experiment 1). As shown in Anderson et al (2000), inhibition is interference dependent. Accordingly, if the elderly did not detect interference, as discussed above, inhibition should not be called into play. This is evidenced not only by the fact that theta power did not decrease across cycles, but also by the absence of a behavioural RIF effect across the two experiments.

Notably, our results are consistent with the Inhibitory Deficit Theory (IDT; Hasher & Zacks, 1988) in that they show impairment in an inhibitory task. Our results advance this theory by identifying a possible reason for this inhibition impairment, which might lie in an earlier stage of interference detection.

Previous research has also pointed in the same direction. For instance, ERP studies in young adults showed that during incongruent trials of a Stroop task (interference inducing trials), a medial frontal negativity (MFN) component occurs between 400 and 500ms (N450; Rebai, Bernard, & Lannou, 1997; West & Alain, 1999), with several studies showing medial prefrontal brain regions to be the generator of this MFN (Hanslmayr et al., 2008; Liotti, Woldorff, Perez, & Mayberg, 2000;West & Alain, 2000). Crucially, the MFN generated by older adults has been shown to be attenuated, during different variants of the Stroop task (West, 2004; West & Schwarb, 2006). Similarly, Tays et al. (2008) found that in a Sternberg-like task, older adults showed a large frontal positivity instead of the MFN, and that this unique pattern of frontal positivity is associated with poorer behavioural performance, rather than with compensatory mechanisms. The fact that a component that is consistently found in interference related trials (such as the MFN) is attenuated in older adults agrees with our results and with the idea that older adults have a harder time detecting interference.

In sum, in the present work, we aimed to understand how age changes the neural dynamics underlying RIF. We aimed to disentangle whether cognitive aging affects interference detection or interference resolution mechanisms, especially in the context of face naming, a task that seems to be rendered especially hard as people age (Lovelace and Twohig, 1990; Maylor, 1990; Naveh-Benjamin et al., 2004). In two experiments we show that the age-related inhibitory deficit largely described in the literature, might be due to a missing early interference signal, with the elderly not detecting interference and consequently not recruiting the inhibitory mechanisms necessary to overcome it. This is, to our knowledge, the first study using electrophysiological measures to understand age-related neural changes underlying RIF and suggesting a specific source for the absence of RIF in the elderly. Going a step further, these findings contribute to the current understanding of the cognitive dynamics during memory retrieval in ageing and how they are reflected in brain oscillations.

## SUPPLEMENTARY FIGURES

**Supplementary Figure 1:**
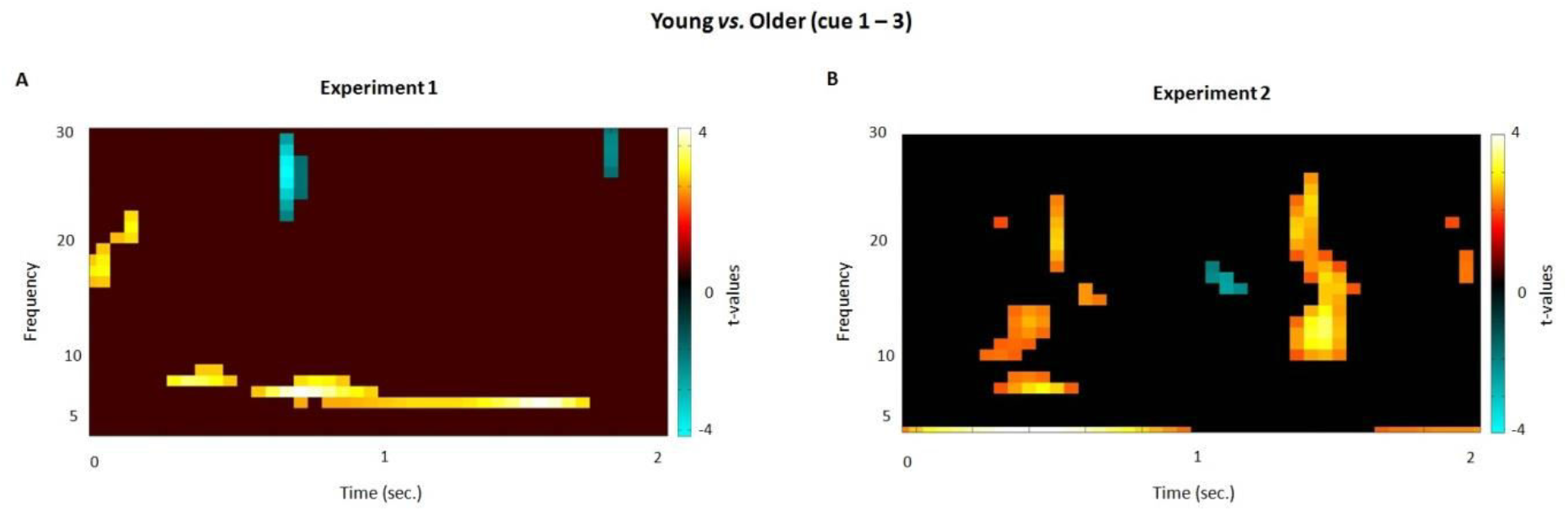
Interaction analysis: time-frequency plots of the differences between younger (cue1 – cue3) and older adults (cue1 – cue3), over all electrodes and for a broader time (0 to 2 sec.) and frequency (0 to 30 Hz) ranges. A) depicts the time-frequency plot obtained from the data of Experiment 1 and B) from Experiment 2.

## Footnotes

1 The sample size was estimated by a power analysis based on the effect size from Ferreira et al., (2014) (*η*^2^_*p*_=0.28); α err prob=0.05 and power=0.95. Estimated sample from the mixed ANOVA resulted in 44 and in the current study was increased up to 48 in order to keep a complete counterbalance of the task material.

